# The RpfB switch is a novel B_12_-sensing riboswitch regulating (non-replicating) persistence in *Mycobacterium tuberculosis*

**DOI:** 10.1101/2024.07.19.603033

**Authors:** Terry Kipkorir, Peter Polgar, Alexandre D’Halluin, Brindha Gap-Gaupool, Vadim A. Makarov, Galina V. Mukamolova, Kristine B. Arnvig

## Abstract

Riboswitches are metabolite-sensing RNA elements that control a wide range of genes in bacteria. Most riboswitches identified to date are broadly conserved and control genes that are directly involved in the transport or biosynthesis of their cognate ligands. However, a minority of switches are restricted to a few species and in addition may bind less obvious ligands. One such switch controls the expression of the *Mycobacterium tuberculosis rpfB* operon, which is critical for resuscitation of dormant bacteria, ribosome maturation and reactivation of latent tuberculosis infection. The switch is restricted to pathogenic mycobacteria and until now, its ligand was unknown. However, in the current study, we identify the ligand as cobalamin or vitamin B_12_. Using in-line probing, we show that vitamin B_12_ binds directly to the riboswitch RNA, and we predict a structure based on the cleavage pattern. Moreover, we show that B_12_ suppresses the expression of an *rpfB-lacZ* reporter fusion and crucially, that B_12_ suppresses resuscitation of *M. tuberculosis* from a state of non-replicating persistence. These findings demonstrate a pivotal role of crosstalk between a host-derived metabolite and a pathogen riboswitch in controlling *M. tuberculosis* persistence with potential for improved interventions.

## Introduction

The ability to sense, respond and ultimately adapt to changing and challenging environments is critical for the survival of all organisms and mediated by a multitude of biomolecules within a cell. Riboswitches are remarkable examples of such molecules. Primarily located in the 5’ leaders of mRNAs they regulate, these RNA elements sense changes in the cellular environment by binding small-molecule ligands and respond with extensive conformational changes that only few proteins can rival. Typically, these ligands are synthesised, transported, or utilised in metabolic pathways involving the genes encoded by the mRNA under riboswitch control [1]. The binding of specific ligands such as ions or metabolites occurs via an aptamer domain, which is often conserved across a wide range of species, suggesting links to early ancestral molecules. However, a few aptamers are only found in a much narrower range of species and therefore potentially more recent in evolutionary terms [2]. Control over the downstream genes is executed by a more variable expression platform, which in most known cases acts through premature termination of transcription/antitermination and/or modulation of ribosome access to the translation initiation region of the mRNA [2-4].

**Mycobacterium tuberculosis,** the cause of human tuberculosis (TB), is exposed to a range of adverse environments during its infection cycle. These include hostile intracellular compartments, and hypoxic, nutrient-deplete extracellular compartments [5]. Each year, millions of people are infected with *M. tuberculosis*, but the majority will control the infection leading to either clearance or development of a latent TB infection (LTBI). Rather than being a single, well-defined condition, LTBI represents a spectrum of disease states with bacteria presumably existing in a dynamic equilibrium between dormancy and active replication [6, 7]. Despite intensive intervention efforts, LTBI remains a major challenge to diagnostics and treatment partly due its potential to progress into active disease at any given time.

**Dormancy** or non-replicating persistence (NRP) is a physiological state of low metabolic activity. It may be induced by antibiotic treatment, nutrient limitation, hypoxia or host defence mechanisms such as nitrosative stress, triggered by nitric oxide (NO), which has been shown to be a potent inducer of NRP *in vitro* (Fig. 1) [8, 9]. Dormant bacteria constitute a heterogeneous population, but commonalities between cells include an enhanced tolerance to antibiotics, host immune assaults, and notably, the dependence on resuscitation promoting factors (Rpfs) for growth [10-12]. *M. tuberculosis* encodes five Rpfs (RpfA-E), which are tightly and differentially regulated on multiple levels to ensure the timely and appropriate response(s) to a variety of stimuli and to prevent excessive degradation of cell-wall peptidoglycan (PG) by the Rpf proteins [4, 13-15]. Hence, it is critical that the cells accurately interpret and respond to their microenvironments. Resuscitation of *M. tuberculosis* and the associated reactivation of disease likely involves a few bacteria sensing the host environment as favourable for growth, followed by the metabolic resuscitation and regrowth of bacteria [14, 16]. What precisely constitutes ‘favourable’ growth conditions and the detailed molecular mechanisms behind dormancy, resuscitation and disease reactivation remain elusive, although host nutrition plays a major role in the ability to control TB infection [17].

**Figure 1:**
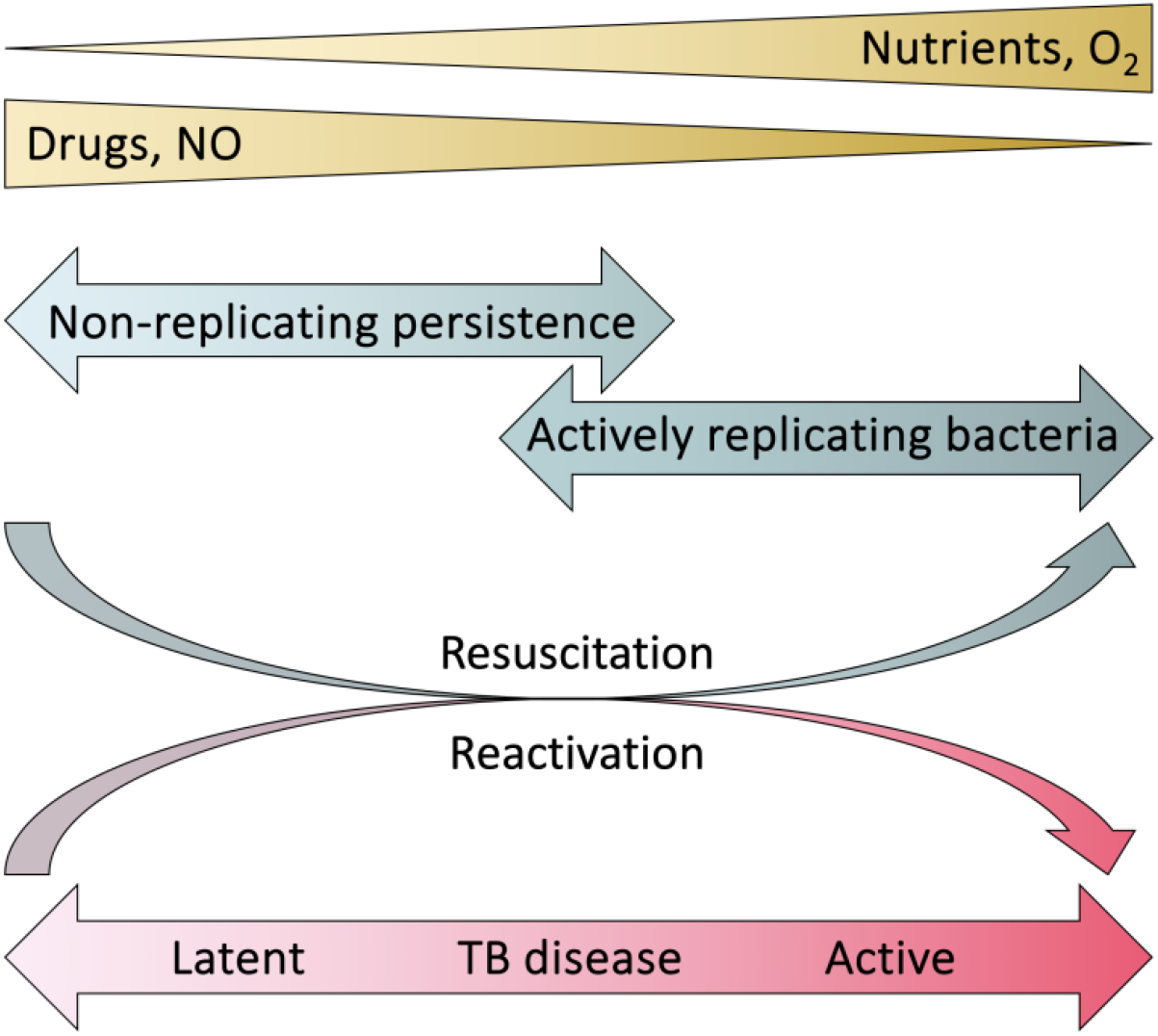
Proposed relationship between NRP, LTBI, bacterial resuscitation and disease reactivation in the context of outside factors that are thought to contribute to these states.

**RpfB** is expressed from the tri-cistronic *rpfB-ksgA-ispE* operon [15]. KsgA is a universally conserved ribosomal RNA (rRNA) methyl transferase required for late stages of ribosomal small subunits (SSU) formation [18]. Deletion of *ksgA* in *Escherichia coli* leads to cold sensitivity, reduced 16S rRNA processing and increased resistance to kasugamycin [18, 19], while interruption of *ksgA* in *M. tuberculosis* by transposon mutagenesis leads to increased sensitivity to clarithromycin [20]. IspE is an essential contributor to the linkage of arabinogalactan to peptidoglycan [21]. Hence, all three gene products of this operon are pivotal in growth-related functions. Notably, despite the presence of multiple Rpfs in *M. tuberculosis*, only RpfB has been directly implicated in reactivation of chronic infection in a mouse model [16].

Expression of the *rpfB* operon is controlled at the transcriptional level by the essential two-component regulator MtrAB [22] and at the post-transcriptional level by an antisense RNA, whose function remains unknown, and a riboswitch whose cognate ligand so far eluded discovery [3, 4, 15]. This riboswitch (the RpfB switch) features two unusual properties: it is the only *M. tuberculosis* riboswitch known to be controlled by intrinsic termination and it has been found in only a subset of pathogenic mycobacteria, making this a rare switch [15, 23].

At the other end of the conservation spectrum, is the clan of B_12_-sensing switches. Identified in thousands of bacteria, this is one of the most widespread families of riboswitches comprised of at least three classes (class I, class IIa and IIb) [2]. Due to structural variations, individual classes recognise variants of vitamin B_12_ or cobalamin, more specifically adenosylcobalamin (AdoB_12_), methylcobalamin (MeB_12_), hydroxocobalamin (HyB_12_) or cyanocobalamin (CNB_12_) [24]. Recently, we characterised the structure-function characteristics of two B_12_-sensing switches from *M. tuberculosis* [25]. The two switches, belonging to class I and II, control the expression of *metE* and *ppe2*, respectively. However, like many other bacteria, *M. tuberculosis* cannot synthesise B_12_ due to the relatively recent deletion of *cobF* and truncation of *cobL* [26, 27]. Therefore, as an intracellular pathogen, *M. tuberculosis* presumably relies entirely on host-derived B_12_, taken up via BacA, suggesting that this ligand could convey information about the host environment in addition to acting as a co-factor in certain metabolic pathways of both host and pathogen [28, 29].

In the current study we identify B_12_ as the cognate ligand for the transcriptional RpfB switch. We predict a secondary structure of the ligand-bound switch and show that the RpfB switch is both smaller and structurally divergent from other known B_12_-sensing switches. Based on this, we propose that the RpfB switch represents a novel class of B_12_ riboswitches. Moreover, we show that B_12_ suppresses downstream *rpfB* expression and importantly, that B_12_ inhibits the resuscitation of *M. tuberculosis* non-replicating persisters. Our findings suggest a role for host-derived vitamin B_12_ as a signal controlling the transition of *M. tuberculosis* persisters to actively replicating cells, thereby providing new insights into host-pathogen interactions with potential consequences for reactivation of LTBI.

## Results

### Overexpression of RpfB switch to identify ligand by sequestration

As the *M. tuberculosis* RpfB switch is not associated with a biosynthetic or transport pathway, deducing its ligand proved challenging. Therefore, we employed an overexpression strategy followed by RNA-seq with the aim of sequestering the unknown ligand and subsequently identifying differentially expressed genes and potentially affected pathways. A replicating plasmid (pKA300) expressing the 130-nucleotide RpfB switch from an anhydrous tetracycline- (ATc)-inducible promoter was transformed into *Mycobacterium smegmatis*, which encodes a homologue of *rpfB* without the riboswitch. The rationale was that a cognate ligand would likely also be present in this background due to the close relationship between the two species, and differential gene expression would be a direct consequence of ligand sequestration due to the absence of this element in *M. smegmatis*. RNA was isolated from log-phase, ATc-induced cultures of *M. smegmatis* and differential gene expression (DEG) analysis performed.

More than 400 genes showed a log2FoldChange (log2FC) ≤ -1 or ≥ 1 (padj ≤0.01; with BH) with 199 up- and 239 downregulated genes. Upregulated genes showed a substantially larger difference in expression with 20 having a log2FC > 6, 38 a log2FC ≥ 5 and representing a range of cellular functions, while the largest downregulation was log2FC = -3 (hypothetical protein, MSMEG_5421, Supplementary table 1). Analysis of all DEG by functional class revealed a striking and highly significant enrichment of genes associated with ribosomal structure and -biogenesis. More than 30 ribosomal proteins and 23S rRNA were significantly downregulated, suggesting a highly specific suppression of translation (Fig. 2 and Supplementary table 1). As no other, single functional group displayed a similar enrichment, the results did not expose a specific, obvious ligand for the RpfB switch, and we proceeded to test a range of metabolites.

**Figure 2:**
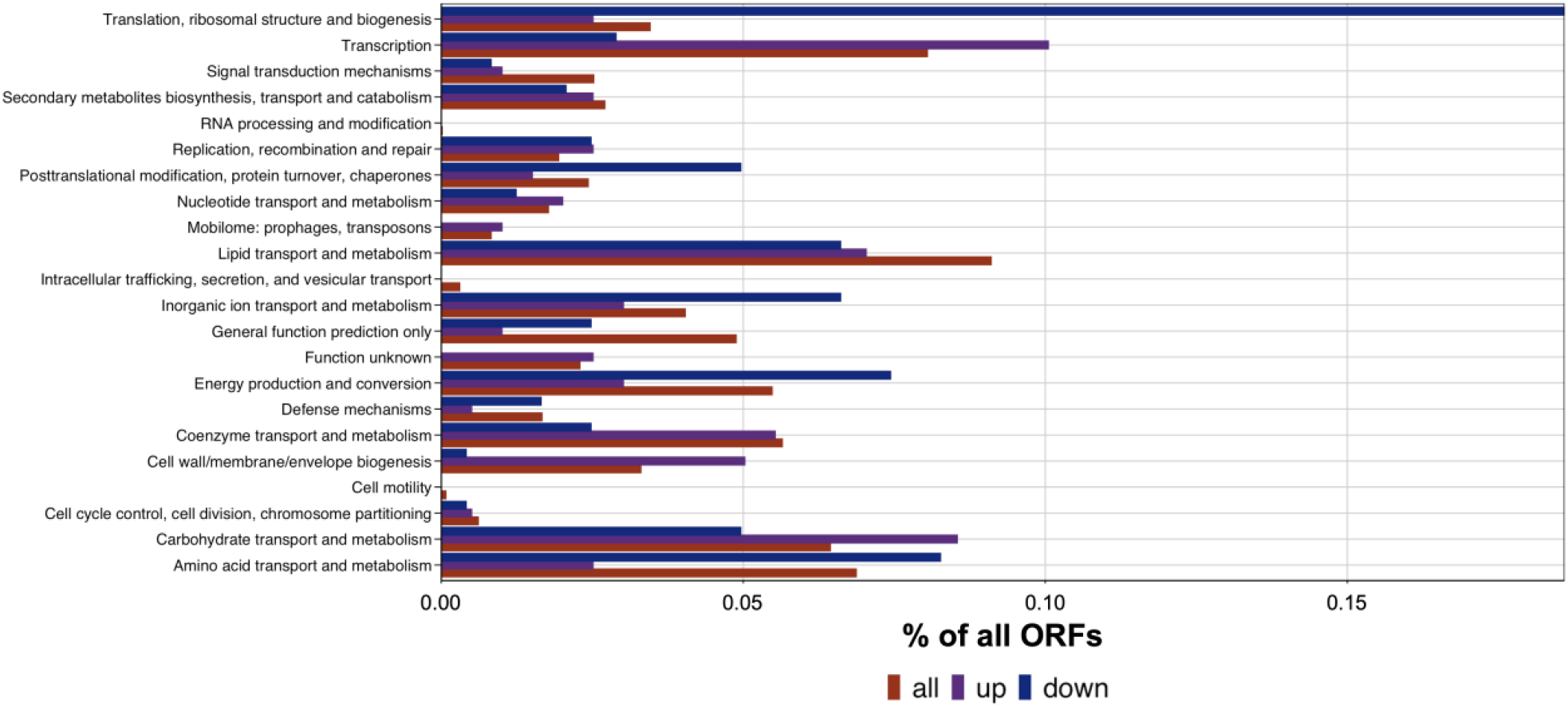
distribution of significantly differentially expressed genes according to functional gene classes. Red indicates frequency of all genes in the *M. tuberculosis* genome, purple indicates genes with Log2Fc ≥1, blue Log2Fc ≤ -1; padj < 0.01. Enrichment of down-regulated genes associated with translation, ribosomal structure and biogenesis (padj = 4.7×10^−21^). The 1670 genes that were not in the COG database have not been included.

### The RpfB Switch represents a novel B_12_-binding element

To probe potential ligands of the switch, a 130-nucleotide ^32^P-5’ labelled transcript of the RpfB switch was generated by in vitro transcription for inline probing and tested against a range of ligands used in our lab including glycine, serine, threonine and CNB_12_.

Neither of the amino acids resulted in any changes in the cleavage pattern, however CNB_12_ resulted in subtle changes in the cleavage pattern in the regions 44-56, especially C53 and 90-130 (Fig. 3A). This led us to test other B_12_ variants, i.e., 1 mM each AdoB_12_, MeB_12_, HyB_12_ and CNB_12_ together with tetrahydrofolate (THF) and *S*-adenosyl-methionine (SAM) associated with B_12_-dependent metabolic pathways and KsgA function. Fig. 3B shows the results of the inline probing, where AdoB_12_ induced a substantial difference in cleavage of the RpfB switch transcript. The other naturally occurring cobalamin, MeB_12_, resulted in slightly fewer changes, while HyB_12_ and CNB_12_ induced only minor changes and THF and SAM had no effect.

**Figure 3:**
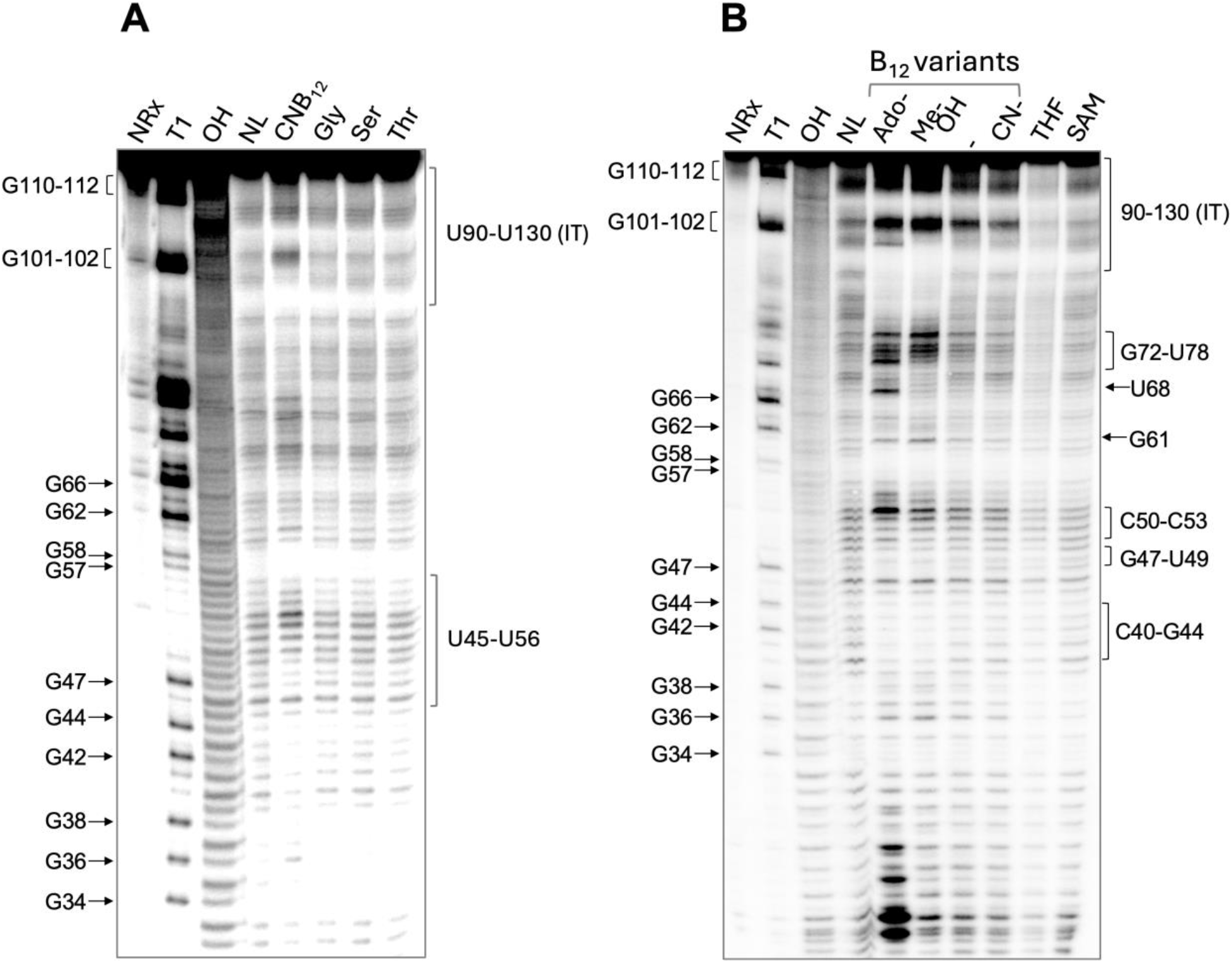
Inline probing of the RpfB switch. Inline suggests that AdoB12 is the preferred ligand for the RpfB switch. A 130-nt in vitro transcribed 5’ 32P-labelled transcript was incubated for 20 hours at 30 °C with 1 mM of the indicated metabolites after which the reactions were separated on a 10% acrylamide gel. Panel A shows reactions with CNB12 and amino acids glycine, serine and threonine; Panel B shows reactions with B_12_ variants and tetrahydrofolate and *S*-adenosyl-methionine.

This strongly suggested that AdoB_12_ is the preferred ligand for the RpfB switch, and we therefore proceeded with further inline probing using a range of AdoB_12_ concentrations to assess binding affinity (Fig. 4). The results indicate that differential cleavage required ligand concentrations > 1mM, suggesting that the affinity between the RpfB switch and AdoB_12_ is relatively low compared to other B_12_-sensing switches in *M. tuberculosis* [25].

**Figure 4:**
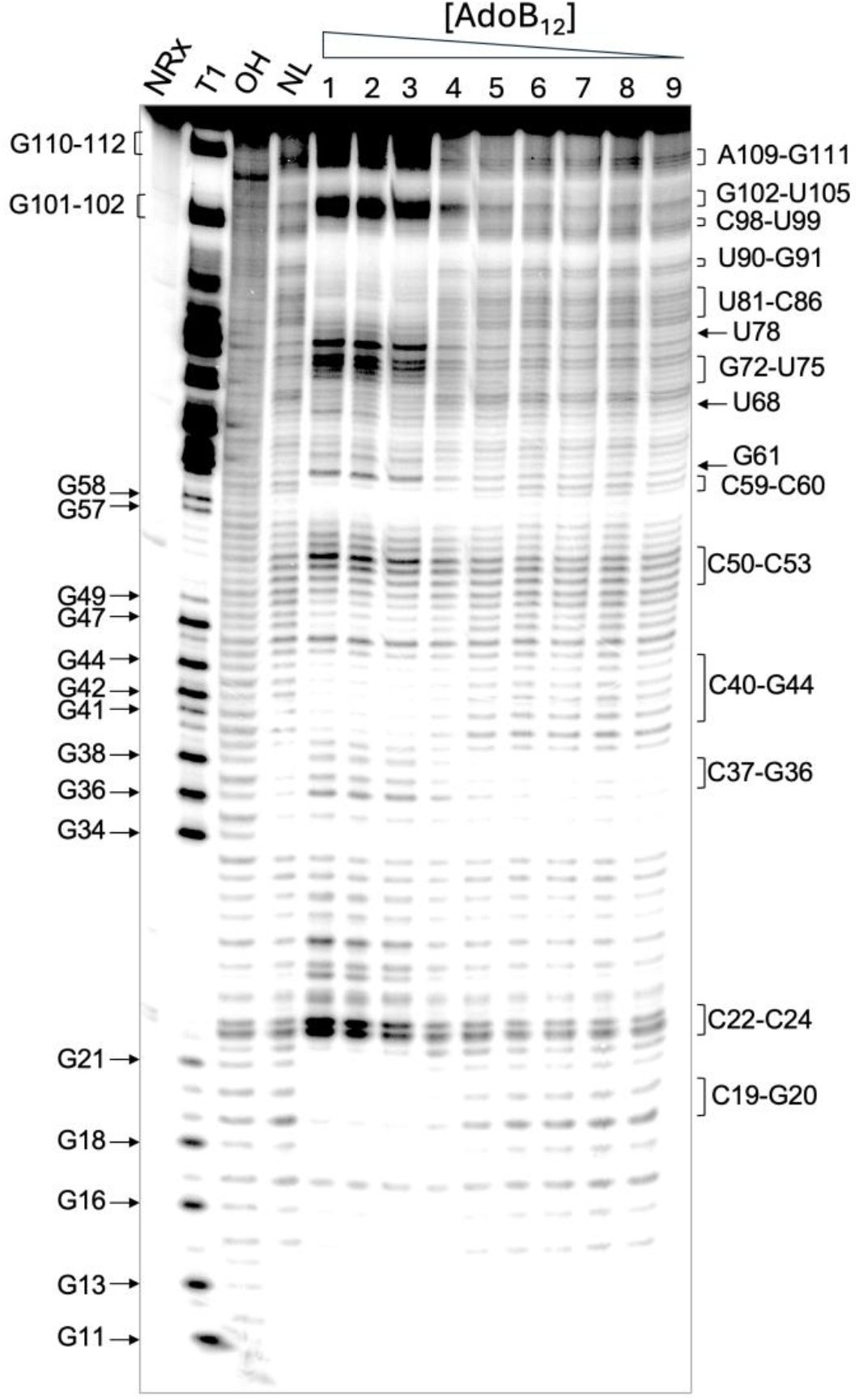
Inline probing of the RpfB Switch using a gradient of AdoB_12_ concentrations: 1 – 4 mM; 2 – 2 mM; 3 – 1 mM; 4 – 100 μM; 5 – 10 μM; 6 – 1 μM; 7 – 100 nM; 8 – 10 nM; 9 – 1 nM. The most pronounced structural modulation occurs with AdoB_12_ concentrations above 1 mM.

Next, we used the cleavage pattern obtained with AdoB_12_ to add constraints to a structure prediction of the switch. The resulting structure, shown in Figure 5, suggests that the B_12_-binding pocket resides in an adenosine-rich, three-way junction, which includes a potential, but non-canonical B_12_ box, shown in green in Fig. 5B. This region is highly conserved between mycobacteria encoding a long RpfB 5’ leader, while the flanking regions show more diversity in species outside the *M. tuberculosis* complex (MTBC) (Fig. 5C). Another typical feature of cobalamin riboswitches is the so-called Kissing Loop (KL), which forms between loops within the aptamer and the expression platform [30]. We identified a potential KL between the first and the last hairpin loops, which would likely stabilise the intrinsic terminator, previously shown to be functional [15].

**Figure 5:**
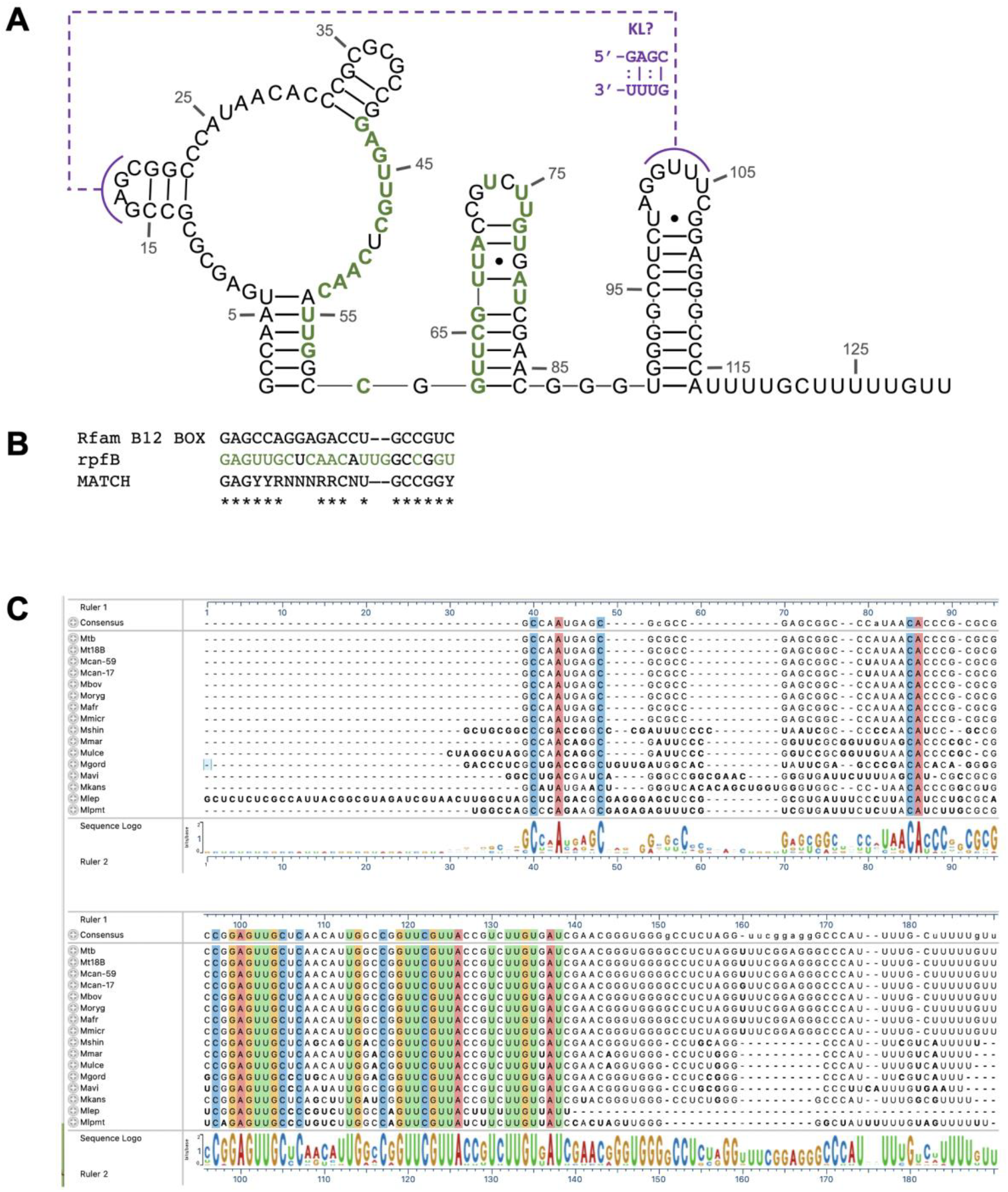
Riboswitch structure and sequence variation. A. Proposed structure of the B_12_-bound RpfB switch. The cleavage pattern obtained in Figs. 3 and 4 were used to add folding constraints to the predicted structure of the switch, which was slightly modified according to regions conserved in mycobacterial elements. This suggests a ligand-binding pocket within a three-way junction. Green residues indicate residues that are highly conserved between mycobacteria with A43 to C53 potentially involved in ligand binding (alignment in Fig. 5C). KL: potential kissing loop interaction between hairpin loops in aptamer and expression platform with basepairing shown in purple. B: Alignment of potential B_12_ box from RpfB with Rfam consensus for the B_12_ box. C: Multiple sequence alignment of *rpfB* leaders from pathogenic mycobacteria using Clustal omega [31]. *Mtb – M. tuberculosis H37Rv; Mt18B – M. tuberculosis 18B; Mcan-59 – M. canettii CIPT 140010059; Mcan-17 - M. canettii CIPT 140070017; Mbov – M. bovis AF2122-97; Moryg – M. orygis 51145; Mafr – M. africanum; Mmicr – M. microti; Mshin – M. shinjukuense; Mmar – M. marinum M; Mulce – M. ulcerans; Mgord – M. gordonae; Mavi – M. avium 104; Mkans – M. kansasii; Mlep – M. leprae TN; Mlpmt – M. lepromatosis Mx1-22A*.

Overall, the results show that the RpfB switch is much smaller and has a visibly different architecture compared to previously described B_12-_sensing switches in *M. tuberculosis* and other bacteria, which all contain a ligand-binding pocket within a *four*-way junction and a highly conserved B_12_ box [24, 25]. Moreover, despite its preference for AdoB_12,_ it does not contain the peripheral extension (P8-P12) characteristic of class I switches [24, 25]. We therefore propose that the RpfB switch represents an entirely novel class (III) of B_12_-sensing riboswitches.

### AdoB_12_ suppresses RpfB expression and resuscitation of *M. tuberculosis* persisters

To investigate the role of B_12_ on *rpfB* expression in vivo, we made a reporter containing the *rpfB* promoter, the RpfB switch and five codons of the RpfB open reading frame fused in frame to *lacZ*. Since Rpfs are generally subject to multi-layered control [15], we also made a reporter in which we had deleted the RpfB switch to ensure that any regulation we observed could be assigned to the switch. To assess the effect of B_12_, both reporters were expressed in *M. smegmatis* wildtype, which is B_12_-producing and a Δ*cobK* mutant, which is B_12_-deficient like *M. tuberculosis*. The resulting β-gal assays, shown in Fig. 6 indicate that the construct without the switch had the same level of expression regardless of the genetic background, i.e. whether there was endogenous B_12_ or not, suggesting that the *rpfB* promoter is not sensitive to the B_12_ status of the cell.

**Figure 6:**
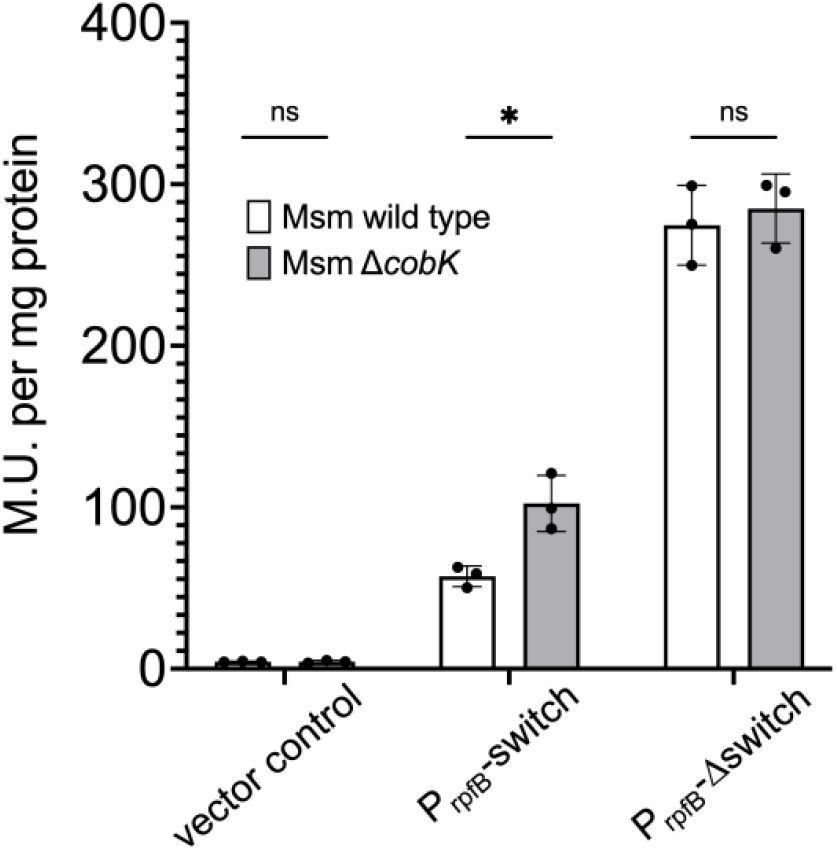
Expression of *rpfB-lacZ* reporters in *Mycobacterium smegmatis*. The bars indicate the β-gal activity of in-frame *rpfB-lacZ* fusions with (P_rpfB_-switch) or without the riboswitch (P_rpfB_-Δswitch) expressed from the *rpfB* promoter in B_12_-producing (wildtype) and B_12_-deficient (Δ*cobK*) backgrounds.

However, the construct that included the switch showed a modest (1.8-fold) but statistically significant increase in the level of β-gal activity in the B_12_-deficient strain, suggesting that the presence of B_12_ led to a reduced expression of RpfB-LacZ via the RpfB switch.

RpfB is believed to be a main contributor to reactivation of LTBI through its resuscitation activity [16]. As the presence of B_12_ suppressed *rpfB* expression, we hypothesised that resuscitation of dormant *M. tuberculosis* might also be suppressed in the presence of B_12_. We used our established NRP-resuscitation model (outlined in Fig. 7A), in which cultures of *M. tuberculosis* were grown to mid-log phase and treated with a custom-made nitric oxide donor to induce differentially culturable NRP cells [8]. Resuscitation was done in 7H9, and the resuscitation index (RI) was calculated (Fig. 7A and methods). To ascertain the effect of AdoB_12_ on the resuscitation of *M. tuberculosis* from NRP, the assay was performed it in the presence or absence of the metabolite.

**Figure 7:**
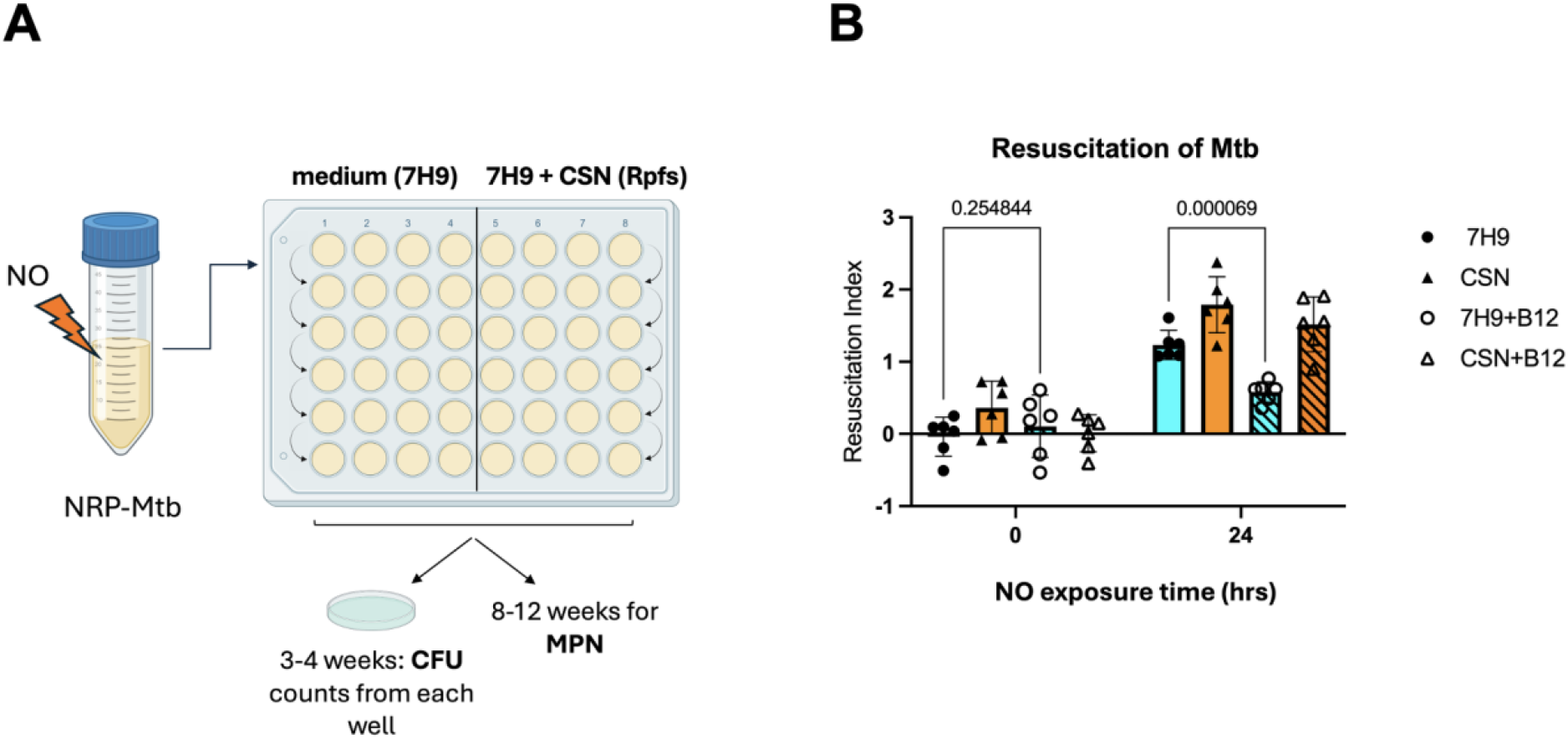
Induction of NRP followed by resuscitation of *M. tuberculosis*. Panel A illustrates the experimental setup for inducing NRP in *M. tuberculosis*. CSN is *M. tuberculosis* culture supernatant, which contains endogenous Rpfs. Panel B shows the results of resuscitation in the presence and absence of AdoB_12_

The results shown in Fig. 7B, indicate a highly significant reduction in the RI between 7H9 and 7H9 + AdoB_12_, while the RI in the presence of Rpf-containing CSN is unaffected. These results suggest that the addition of AdoB_12_ suppresses *M. tuberculosis* resuscitation from NRP in an RpfB-dependent manner, which is likely to have consequences for the activation of LTBI.

Together our results strongly suggest that vitamin B_12_, in particular AdoB_12_, likely obtained from the host, controls the expression of the *rpfB* operon and more importantly, inhibits resuscitation from NRP/dormancy.

## Discussion

Understanding the conditions that drive the transition between active growth and NRP in *M. tuberculosis* remain one of the keys to designing new intervention strategies against active disease as well as LTBI. In the current study, we reveal a piece of this puzzle in the form of a new B_12_-sensing riboswitch that controls the expression of *M. tuberculosis* RpfB, and thus resuscitation of non-replicating persisters is suppressed.

Based on its size, predicted structure and the absence of elements conserved in other B_12_-sensing switches, we propose that the RpfB switch represents a new class of B_12_ switches, which to our knowledge is only found in a subset of pathogenic mycobacteria [15] and Fig. 5. We currently do not know whether switch variants encoded by pathogenic but non-tuberculous mycobacteria (NTM) is functional. High expression of the switch in *M. smegmatis* did not immediately point to B_12_ as the ligand, since a key indicator would presumably be upregulation of the B_12_-regulated *metE*, which was in fact downregulated. It is possible that an induction time of two hours before RNA isolation enabled downstream/secondary effects, which obscured the results. We did however, observe an upregulation of some genes involved in cobalamin biosynthesis, e.g. *cobD,G,L,T* (Supplementary table 1).

The central region of the switch, from G42 to C82, is highly conserved in all the mycobacteria we aligned, while the flanking regions showed substantial variations in species outside the MTBC. With respect to known classes of B_12_ switches, there were limited similarities. In the RpfB switch, AdoB_12_ appears to bind within a central purine-rich loop, which forms a three-way junction, as opposed to a four-way junction in previously characterised B_12_ switches. The first section of the mycobacterium-conserved region mentioned above formed part of this loop (G42-C53) and showed some similarity to the B_12_ box critical for ligand binding in canonical B_12_ switches. Less conserved elements included a potential KL that would likely stabilise the intrinsic terminator responsible for controlling the genetic output of the operon. The poor conservation of elements deemed critical for ligand binding in other classes of B_12_-sensing switches may in part explain the relatively low affinity of the switch to its primary ligand, AdoB_12_. In addition to the low affinity, this switch functions via a transcriptional expression platform, which is likely to require higher levels of ligand for control [32, 33]. This raises the question of when, during infection, and in which micro-environment B_12_ levels are sufficient to suppress *rpfB* expression. Vitamin B_12_ remains tightly bound to proteins in every cellular compartment except for the lysosome. Here the cofactor is stripped from transcobalamin before being rebound for export by lysosomal transporters [34]. Thus, the lysosome is presumably the only host compartment, where *M. tuberculosis* encounters free B_12_. This in turn suggests that only the subpopulation of bacteria that do not escape the phagosome nor prevent phago-lysosome fusion [35], will encounter the B_12_ required for suppression of resuscitation. It has recently been shown that a lysosome-poor subset of immune cells harbours ∼5-fold more live *M. tuberculosis* than their lysosome-rich counterparts [36]. Future investigations should reveal if transient occupation of a lysosome drives NRP via the RpfB switch, leading to a lower number of *M. tuberculosis* cells.

Often, riboswitches control genetic outputs in a coordinated fashion within an organism [1]. However, the role of vitamin B_12_ and its cognate switches in *M. tuberculosis* growth and pathogenesis remains elusive and confusing. Since *M. tuberculosis* cannot synthesise B_12_, the presence/levels of this metabolite in the bacilli are host-derived and hence could serve to inform about favourable versus non-favourable growth conditions. In mammals, B_12_ deficiency affects innate and adaptive immune functions. Experiments in B_12-_deficient rats and mice indicate a reduction in the levels of Serum C3, IgM, IgG and CD4+IFN-gamma+ cells [37, 38], which is mirrored by the finding that human B_12_ deficiency correlates with diminished immune functions including impaired NK cell activity and reduced levels of C3, C4, Ig, CD4+ and CD8+ T cells, all of which can be restored by B_12_ supplements [39, 40]. Moreover, it was shown that B_12_-deficient lambs have a significantly diminished cell-mediated immune response to *Mycobacterium paratuberculosis* [41]. Finally, a recent study showed that *M. tuberculosis* exhibits reduced virulence in B_12_-deficient mice compared to mice fed a B_12_ replete diet [42]. Together, these findings suggest that B_12_ deficiency has negative consequences for the ability to control infections including TB.

We have shown that B_12_ suppresses resuscitation via the RpfB switch and the simultaneous expression of PPE2 [25], which otherwise suppresses the production of host iNOS [43]. Based on this, high B_12_ levels may favour higher NO production, leading to a more hostile host environment and prolonged persistence. This is supported by a study showing that the B_12_-transporter, BacA may be required for maintaining a chronic infection in mice [44].

On the other hand, B_12_ suppresses the expression of *metE* [25, 45], while enabling the activity of the more efficient isozyme, MetH. This could be associated with enhanced growth, in turn suggesting a less hostile environment. In this context it is also worth noting that a transposon screen for B_12_ ‘sensitivity’ in a Δ*metH* strain of *M. tuberculosis*, resulted in a single insertion in *rpfB* [28]. We have so far not been able to explain this result and how it might be connected to the RpfB switch.

It remains to be seen whether enhancing NRP represents an advantage for patients by suppression of active disease. It is possible that NRP facilitates immune-mediated clearance, but it could also obscure antibiotic-mediated killing of the pathogen.

A recently published study provided strong evidence that nutrition plays a critical role in controlling TB disease [17]. Our study suggests a molecular mechanism by which vitamin B_12_ could play a role in such control. Perhaps paying heed to B_12_ serum levels of patients and potentially adding supplements could enhance drug treatment? Moreover, in a world with decreasing food security and more people turning to vegan diets, the role of B_12_ needs to be fully understood to prevent potentially increased susceptibility to TB and other infections in the global population.

## Materials and methods

### Strains, plasmids, cloning, growth

In-frame *lacZ* fusions were made in pKA425 described in [15] by inserting gene blocks between the HindIII and NcoI sites using Gibson assembly. The plasmids harboured a region that included the *rpfB* promoter and 5 codons of the *rpfB* coding region with or without the switch in between. Plasmids, including the empty vector were transformed into *M. smegmatis MC2 155* or △*cobK* (provided by Professor Digby Wagner, University of Cape Town) by electroporation.

For RNA-seq, cultures of *M. smegmatis* were grown to OD600∼0.4 before expression of the RpfB switch was induced by adding 500 ng/ml anhydrotetracycline (ATc) to the cultures and samples were removed for RNA isolation 2 hours post induction.

*M. tuberculosis* H37Rv strain was provided by Professor W.R. Jacobs (Albert Einstein College of Medicine). Mycobacteria were grown in Middlebrook 7H9 broth, supplemented with 0.2% (v/v) glycerol, 10% (v/v) albumin-dextrose-catalase (ADC), and 0.05% (w/v) Tween 80 (hereafter 7H9 medium). Culture supernatants (CSN) were prepared from exponentially growing cultures (OD_580nm_ of 0.6-0.8) and sterilized by double filtration (0.22 μm filters). AdoB_12_ was obtained from Thermo Scientific.

### RNA isolation and purification

RNA was isolated as previously described [25]. Briefly, cells were cold shocked with 30% ice, pelleted and total RNA was extracted using the FastRNA Pro Blue Kit (MP Biomedicals). The samples were treated with the Turbo DNase (ThermoFisher) until no signal was detected by PCR using the RedTaq readymix (Sigma). RNA concentration and purity were assessed on a Nanodrop 2000 and RNA integrity was checked by gel electrophoresis.

### RNA-seq and data analysis

RNA was treated with DNase until PCR clean. Samples were sequenced by the sequencing facility at UCL Genomics, Institute of Child Health. Obtained sequences were mapped against the *M. smegmatis* genome (NCBI RefSeq) with Bowtie2 [46], and the resulting SAM file converted to binary BAM, sorted and indexed with SAMtools [47]. Mapped reads were assigned to genes using featureCounts [48] and the resulting matrix directly fed into DESeq2 [49] for differential gene expression analysis. GO analysis was performed using custom R code (https://github.com/ppolg/rpfB) according to Database of Clusters of Orthologous Genes -COG), https://www.ncbi.nlm.nih.gov/research/cog). Sequencing data are available at ArrayExpress, accession number E-MTAB-14260.

### Probing and structure determination

The structure probing for the RpfB switch was carried out as previously described [25]. Briefly, a 147-nucleotide amplicon was generated by PCR from *M. tuberculosis* genomic DNA. A 130-nucleotide transcript was in-vitro transcribed using the PCR amplicon as template and purified from a denaturing 8% polyacrylamide gel using the crush-and-soak method (0.5 mM NaOAc, pH5.2, 1mM EDTA, pH8.0, 0.1% SDS, 30 μl acidic phenol). For probing reactions, 10 pmol of 5’ end-labelled RNA was mixed with inline buffer and varying concentrations of ligands and incubated at 30 ^°^C for 20 hours. B_12_ variants were from Thermo Scientific; all other metabolites were purchased from Sigma. Cleavage reactions were resolved on 8% polyacrylamide sequencing gel, and subsequently visualised and analysed on a Typhoon FLA 9500 Phosphorimager (GE).

### Reporter assays

Three single colonies from each successful transformation, representing biological replicates, were grown in 30 mL 7H9/ADC media until mid-log phase and cell pellets were harvested by centrifugation. The β-gal activity was determined as previously described [25] normalising to total protein content using ThermoFisher’s BCA kit following the manufacturer’s recommendations. The data were analysed using Graphpad Prism (version 10).

### Resuscitation assays

Cultures of *M. tuberculosis* were grown to mid-log phase and treated with a custom-made nitric oxide donor 3-cyano-5-nitropyridin-2-yl diethyldithiocarbamate (100 μM) for 24 hours at 37°C statically to induce differentially culturable NRP [8]. Treated *M. tuberculosis* were subsequently serially diluted in 48-well microplates by adding 50 μl of cells to 450 μl of resuscitation medium either standard 7H9 or 7H9 with Rpf-containing CSN, with or without 1 mM AdoB_12_ (Thermo Scientific) for most probable number (MPN) determination. For each condition 4 replicate wells were used to calculate an MPN count for one biological sample using the MPN calculator [50].

For CFU counts, 10 μl from each well corresponding to 10_-1_-10_-5_ dilutions from the MPN plates (7H9 and CSN) were spotted onto Middlebrook 7H10 agar plates. MPN and CFU plates were sealed and incubated at 37°C for up to 12 weeks. The Resuscitation Index (RI) was calculated as RI=Log_10_ MPN/ml - Log_10_ CFU/ml.

## Acknowledgments

The authors thank Digby Warner for supplying *M. smegmatis* Δ*cobK* and William Jacobs for *M. tuberculosis* H37Rv.

## Author Contributions

TK, KBA, GVM designed the study. TK, AD, BGG, PP, GVM, KBA conducted experiments. TK, AD, PP, GVM, KBA performed data analysis and wrote the manuscript. VAM provided reagents.

## References

1. Breaker, R.R., Prospects for riboswitch discovery and analysis. Mol Cell, 2011. 43(6): p. 867–79.

2. McCown, P.J., et al., Riboswitch diversity and distribution. RNA, 2017. 23(7): p. 995–1011.

3. Arnvig, K.B., Riboswitches: choosing the best platform. Biochem Soc Trans, 2019. 47(4): p. 1091–1099.

4. Schwenk, S. and K.B. Arnvig, Regulatory RNA in Mycobacterium tuberculosis, back to basics. Pathog Dis, 2018. 76(4).

5. Cambier, C.J., S. Falkow, and L. Ramakrishnan, Host evasion and exploitation schemes of Mycobacterium tuberculosis. Cell, 2014. 159(7): p. 1497–509.

6. Barry, C.E., 3rd, et al., The spectrum of latent tuberculosis: rethinking the biology and intervention strategies. Nat Rev Microbiol, 2009. 7(12): p. 845–55.

7. Dartois, V.A. and E.J. Rubin, Anti-tuberculosis treatment strategies and drug development: challenges and priorities. Nat Rev Microbiol, 2022. 20(11): p. 685–701.

8. Glenn, S., et al., Exposure to nitric oxide drives transition to differential culturability in Mycobacterium tuberculosis. bioRxiv, 2023.

9. Voskuil, M.I., et al., Inhibition of respiration by nitric oxide induces a Mycobacterium tuberculosis dormancy program. J Exp Med, 2003. 198(5): p. 705–13.

10. Mukamolova, G.V., et al., Resuscitation-promoting factors reveal an occult population of tubercle Bacilli in Sputum. Am J Respir Crit Care Med, 2010. 181(2): p. 174–80.

11. Turapov, O., et al., The in vivo environment accelerates generation of resuscitation-promoting factor-dependent mycobacteria. Am J Respir Crit Care Med, 2014. 190(12): p. 1455–7.

12. Turapov, O., et al., Phenotypically Adapted Mycobacterium tuberculosis Populations from Sputum Are Tolerant to First-Line Drugs. Antimicrob Agents Chemother, 2016. 60(4): p. 2476–83.

13. Gupta, R.K., B.S. Srivastava, and R. Srivastava, Comparative expression analysis of rpf-like genes of Mycobacterium tuberculosis H37Rv under different physiological stress and growth conditions. Microbiology (Reading), 2010. 156(Pt 9): p. 2714–2722.

14. Rosser, A., et al., Resuscitation-promoting factors are important determinants of the pathophysiology in Mycobacterium tuberculosis infection. Crit Rev Microbiol, 2017. 43(5): p. 621–630.

15. Schwenk, S., et al., Cell-wall synthesis and ribosome maturation are co-regulated by an RNA switch in Mycobacterium tuberculosis. Nucleic Acids Res, 2018. 46(11): p. 5837–5849.

16. Tufariello, J.M., et al., Deletion of the Mycobacterium tuberculosis resuscitation-promoting factor Rv1009 gene results in delayed reactivation from chronic tuberculosis. Infect Immun, 2006. 74(5): p. 2985–95.

17. Bhargava, A., et al., Nutritional supplementation to prevent tuberculosis incidence in household contacts of patients with pulmonary tuberculosis in India (RATIONS): a field-based, open-label, cluster-randomised, controlled trial. Lancet, 2023. 402(10402): p. 627–640.

18. Maksimova, E., et al., Protein Assistants of Small Ribosomal Subunit Biogenesis in Bacteria. Microorganisms, 2022. 10(4).

19. Connolly, K., J.P. Rife, and G. Culver, Mechanistic insight into the ribosome biogenesis functions of the ancient protein KsgA. Mol Microbiol, 2008. 70(5): p. 1062–75.

20. Phunpruch, S., et al., A role for 16S rRNA dimethyltransferase (ksgA) in intrinsic clarithromycin resistance in Mycobacterium tuberculosis. Int J Antimicrob Agents, 2013. 41(6): p. 548–51.

21. Eoh, H., P.J. Brennan, and D.C. Crick, The Mycobacterium tuberculosis MEP (2C-methyl-d-erythritol 4-phosphate) pathway as a new drug target. Tuberculosis (Edinb), 2009. 89(1): p. 1–11.

22. Sharma, A.K., et al., MtrA, an essential response regulator of the MtrAB two-component system, regulates the transcription of resuscitation-promoting factor B of Mycobacterium tuberculosis. Microbiology (Reading), 2015. 161(6): p. 1271–81.

23. D’Halluin, A., et al., Premature termination of transcription is shaped by Rho and translated uORFS in Mycobacterium tuberculosis. iScience, 2023. 26(4): p. 106465.

24. Polaski, J.T., et al., Cobalamin riboswitches exhibit a broad range of ability to discriminate between methylcobalamin and adenosylcobalamin. J Biol Chem, 2017. 292(28): p. 11650–11658.

25. Kipkorir, T., et al., A novel regulatory interplay between atypical B12 riboswitches and uORF translation in Mycobacterium tuberculosis. Nucleic Acids Res, 2024.

26. Ngabonziza, J.C.S., et al., A sister lineage of the Mycobacterium tuberculosis complex discovered in the African Great Lakes region. Nat Commun, 2020. 11(1): p. 2917.

27. Supply, P., et al., Genomic analysis of smooth tubercle bacilli provides insights into ancestry and pathoadaptation of Mycobacterium tuberculosis. Nat Genet, 2013. 45(2): p. 172–9.

28. Gopinath, K., et al., A vitamin B(1)(2) transporter in Mycobacterium tuberculosis. Open Biol, 2013. 3(2): p. 120175.

29. Mulholland, C.V., et al., Propionate prevents loss of the PDIM virulence lipid in Mycobacterium tuberculosis. Nat Microbiol, 2024.

30. Johnson, J.E., Jr., et al., B12 cofactors directly stabilize an mRNA regulatory switch. Nature, 2012. 492(7427): p. 133–7.

31. Madeira, F., et al., The EMBL-EBI Job Dispatcher sequence analysis tools framework in 2024. Nucleic Acids Res, 2024. 52(W1): p. W521–W525.

32. Wickiser, J.K., et al., The kinetics of ligand binding by an adenine-sensing riboswitch. Biochemistry, 2005. 44(40): p. 13404–14.

33. Wickiser, J.K., et al., The speed of RNA transcription and metabolite binding kinetics operate an FMN riboswitch. Mol Cell, 2005. 18(1): p. 49–60.

34. Hannibal, L. and D.W. Jacobsen, Intracellular processing of vitamin B(12) by MMACHC (CblC). Vitam Horm, 2022. 119: p. 275–298.

35. VanderVen, B.C., et al., The Minimal Unit of Infection: Mycobacterium tuberculosis in the Macrophage. Microbiol Spectr, 2016. 4(6).

36. Zheng, W., et al., Mycobacterium tuberculosis resides in lysosome-poor monocyte-derived lung cells during chronic infection. PLoS Pathog, 2024. 20(5): p. e1012205.

37. Funada, U., et al., Vitamin B-12-deficiency affects immunoglobulin production and cytokine levels in mice. Int J Vitam Nutr Res, 2001. 71(1): p. 60–5.

38. Funada, U., et al., Changes in CD4+CD8-/CD4-CD8+ ratio and humoral immune functions in vitamin B12-deficient rats. Int J Vitam Nutr Res, 2000. 70(4): p. 167–71.

39. Erkurt, M.A., et al., Effects of cyanocobalamin on immunity in patients with pernicious anemia. Med Princ Pract, 2008. 17(2): p. 131–5.

40. Tamura, J., et al., Immunomodulation by vitamin B12: augmentation of CD8+ T lymphocytes and natural killer (NK) cell activity in vitamin B12-deficient patients by methyl-B12 treatment. Clin Exp Immunol, 1999. 116(1): p. 28–32.

41. Vellema, P., et al., The effect of cobalt supplementation on the immune response in vitamin B12 deficient Texel lambs. Vet Immunol Immunopathol, 1996. 55(1-3): p. 151–61.

42. Campos-Pardos, E., et al., Dependency on host vitamin B12 has shaped Mycobacterium tuberculosis Complex evolution. Nat Commun, 2024. 15(1): p. 2161.

43. Bhat, K.H., et al., The PPE2 protein of Mycobacterium tuberculosis translocates to host nucleus and inhibits nitric oxide production. Sci Rep, 2017. 7: p. 39706.

44. Domenech, P., et al., BacA, an ABC transporter involved in maintenance of chronic murine infections with Mycobacterium tuberculosis. J Bacteriol, 2009. 191(2): p. 477–85.

45. Warner, D.F., et al., A riboswitch regulates expression of the coenzyme B12-independent methionine synthase in Mycobacterium tuberculosis: implications for differential methionine synthase function in strains H37Rv and CDC1551. J Bacteriol, 2007. 189(9): p. 3655–9.

46. Langmead, B. and S.L. Salzberg, Fast gapped-read alignment with Bowtie 2. Nat Methods, 2012. 9(4): p. 357–9.

47. Li, H., et al., The Sequence Alignment/Map format and SAMtools. Bioinformatics, 2009. 25(16): p. 2078–9.

48. Liao, Y., G.K. Smyth, and W. Shi, featureCounts: an efficient general purpose program for assigning sequence reads to genomic features. Bioinformatics, 2014. 30(7): p. 923–30.

49. Love, M.I., W. Huber, and S. Anders, Moderated estimation of fold change and dispersion for RNA-seq data with DESeq2. Genome Biol, 2014. 15(12): p. 550.

50. Jarvis, B., C. Wilrich, and P.T. Wilrich, Reconsideration of the derivation of Most Probable Numbers, their standard deviations, confidence bounds and rarity values. J Appl Microbiol, 2010. 109(5): p. 1660–7.

